## Supplementary Material for "The potential of gene drive releases in malaria vector species to reduce the malaria burden in different African environments"

#### Table of Contents

S1. Sources of geospatial data used in models of gene drive dynamics and impacts on malaria ... 3
S2. Variables used in selecting representative environments ... 8
S3. Mosquito metapopulation model ... 8
S3.1 Overview ... 8
S3.2 Modelling the landscape ... 9
3.2.1 Settlement locations ... 9
3.2.2 Settlement localities ... 9
3.2.3 Human population per settlement ... 9
3.2.4 Water courses ... 9
3.2.5 Rainfall ... 10
3.2.6 Species distributions ... 10
S3.3 Local population dynamics ... 10
3.3.1 Juvenile stage ... 10
3.3.2 Setting the area-specific carrying capacities ... 11
3.3.3 Adult stage ... 12
3.3.4 Dispersal ... 12
3.3.5 Gene drive ... 13
3.3.6 Parameters of models of mosquito and gene drive dynamics ... 13
S4. Modelling malaria infection dynamics in humans ... 17
S4.1 Malaria infection dynamics in mosquitoes ... 25
S4.2 Estimating vector abundances ... 26
S4.3 Treatment and Seasonal Malaria Chemoprevention ... 26
S4.4 Insecticide-treated bednets ... 27
4.4.1 Impacts on vector mortality and blood-feeding ... 27
4.4.2 Bednet distributions ... 29

### S1. Sources of geospatial data used in models of gene drive dynamics and impacts on malaria

Sources of geospatial data are provided in Table S1 and the methods used to aggregate fine resolution geospatial data to produce area-level values are described in Table S2.

**Table S1.** Description of geospatial data sources used in mosquito population and malaria transmission dynamic models

| Short name | Description | Temporal resolution | Spatial resolution | Data year | URL | Date accessed | Citation |
|---|---|---|---|---|---|---|---|
| *Plasmodium falciparum* prevalence | | | | | | | |
| Pf prevalence | Proportion of Children 2 to 10 years of age showing, on a given year, detectable *Plasmodium falciparum* parasite | Annual | 5km | 2019 | https://data.malariaatlas.org/maps?layers=Malaria:202206_Global_Pf_Parasite_Rate | 01/06/2023 | [1] |
| Vector complex composition | | | | | | | |
| Arabiensis vs gambiae/coluzzii | Proportional abundance of *An. arabiensis* relative to that of *An. gambiae* and *An. coluzzii* combined | Static | 2.5 arcminutes (approx. 5km) | N/A | No URL (sourced from author) | N/A | [2] |
| Funestus vs An. gambiae complex species | Proportional abundance of *An. funestus* relative to that of *An. gambiae, An. coluzzii* and *An. arabiensis* combined | Static | 2.5 arcminutes (approx. 5km) | N/A | No URL (sourced from author) | N/A | [2] |
| Landscape variables | | | | | | | |
| Settlement locations | The presence or absence of buildings ('World Settlement Footprint - Sentinel-1/Sentinel-2 - Global, 2019') | Annual | 10m | 2019 | https://download.geoservice.dlr.de/WSF2019/files/ | 01/06/2023 | [3] |
| Human population | WorldPop human distribution | Annual | 1km | 2020 | https://hub.worldpop.org/geodata/listing?id=64 | 01/06/2023 | [4] |

| | | | | | | | |
|---|---|---|---|---|---|---|---|
| Watercourses | Rivers and lakes | Static | N/A (vector format) | N/A | https://www.hydrosheds.org/hydroatlas | 01/06/2023 | [5] [6] |
| Total precipitation | Accumulated water that falls to the Earth's surface (m) | Hourly | 0.25 degrees | 1979-2021 | https://cds.climate.copernicus.eu/cdsapp#!/dataset/reanalysis-era5-single-levels?tab=overview | 01/06/2023 | [7] |
| Drug treatment, SMC, vector control | | | | | | | |
| Treatment seeking | The population at risk weighted mean treatment coverage of an effective antimalarial for each site are taken from the malaria atlas project raster entitled: “Effective treatment with an Antimalarial drug version 2020” | Annual | Admin-1 | 2000-2018 | https://data.malariaatlas.org/maps?layers=Interventions:202106_Global_Antimalarial_Effective_Treatment,Malaria:202206_Global_Pf_Parasite_Rate | 01/01/2020 | [8-10] |
| ACT treatment coverage | Proportion of clinical cases that seek treatment that are given ACT | Annual | Admin-1 | 2000-2018 | https://dhsprogram.com/ | 01/01/2020 | [8-10] |
| SMC coverage | Proportion of 2-5 year olds receiving annual Seasonal Malaria Chemoprevention (SP-AQ) | Annual | Admin-1 | 2000-2018 | https://www.smc-alliance.org/ | 01/01/2020 | [10] SMC Alliance, (2023) |
| ITN coverage | Proportion of population that sleeps under an ITN during a defined year | Annual | 5km | 2000-2018 | https://data.malariaatlas.org/maps?layers=Interventions:202106_Africa_Insecticide_Treated_Net_Access,Malaria:202206_Global_Pf_Parasite_Rate | 01/06/2023 | [11] |
| Coverage of IRS by insecticide class | Coverage of indoor residual spraying with insecticide (proportion of households sprayed) | Annual | Admin-0 | 2000-2018 Values for the year 2018 were estimated | https://figshare.com/articles/dataset/Indoor_residual_spraying_IRS_coverage_data_from_1997-2017/10066910/1 | 01/06/2023 | [12] |

| | | | | | | | |
|---|---|---|---|---|---|---|---|
| | | | | from the data layer for the year 2017. | | | |
| Insecticide resistance of vectors | Phenotypic resistance to insecticides in mosquito species from the An. gambiae complex in Sub-Saharan Africa | Annual | 2.5 arcminutes (approx. 5Km) | 2005-2017 Values for years 2000-2004 were estimated from the data layer for 2005. Values for the year 2018 were estimated from the data layer for the year 2017. | https://figshare.com/articles/dataset/Mapping_trends_in_insecticide_resistance_phenotypes_in_African_malaria_vectors/9912623 | 01/06/2023 | [13] |

**Table S2.** Methodology for processing geospatial data layers to produce aggregate measures for 1 x 1 degree area

| **Short name** | **Aggregation function** | **Aggregate measure per 1 x 1 degree area** | **Use in study** |
|---|---|---|---|
| Rainfall | | | |
| Total precipitation | Mean | Average weekly rainfall (mm) within the time period January 1st 1979 to December 31st 2021 at all points on the lattice within our study area (25 points per one- degree grid square). | Area selection |
| Max dry weeks | Maximum | The maximum of the average number of dry weeks (averaged over years) across all the locations in the square. The threshold for a dry week was less than one mm per week. | Area selection |
| *Plasmodium falciparum* prevalence | | | |
| Pf prevalence | Mean | Calculated by multiplying together the fine resolution layers for *P. falciparum* prevalence and population size, then summing the quantities across all values (pixels) within each area, and dividing the result by the total population size in each square. | Area selection **and** Parameterising models of gene drive dynamics in mosquitoes and malaria transmission in humans |
| Vector complex composition | | | |
| Proportion *An. arabiensis* | Mean | The average over the area of the abundance of *An. arabiensis* relative to *An. gambiae* and *An. coluzzi* combined | Area selection **and** Parameterising models of gene drive dynamics in mosquitoes and malaria transmission in humans |
| Proportion *An. funestus* | Mean | The average over the area of the abundance of *An. funestus* relative to *An. arabiensis, An. gambiae* and *An. coluzzi* combined | Area selection **and** Parameterising models of gene drive dynamics in mosquitoes and malaria transmission in humans |
| Drug treatment, SMC, vector control | | | |

| | | | |
|---|---|---|---|
| Treatment seeking | Mean weighted by the proportion of each Admin1 unit that intersects each area | The population at risk weighted mean treatment coverage of an effective antimalarial | Parameterising model of malaria transmission in humans |
| ACT drug treatment coverage | Mean weighted by the proportion of each Admin1 unit that intersects each area | Proportion of clinical cases that seek treatment that are given ACT | Parameterising model of malaria transmission in humans |
| SMC coverage | Mean weighted by the proportion of each Admin1 unit that intersects each area | Proportion of 2-5 year olds receiving annual Seasonal Malaria Chemoprevention (SP-AQ) | Parameterising model of malaria transmission in humans |
| ITN coverage | Mean weighted by human population size in each pixel | Proportion of population that sleeps under an ITN during a defined year | Parameterising models of gene drive dynamics in mosquitoes and malaria transmission in humans |
| Coverage of IRS by insecticide class | Mean weighted by human population size in each pixel | Coverage of indoor residual spraying with insecticide (proportion of households sprayed) | Parameterising models of gene drive dynamics in mosquitoes and malaria transmission in humans |
| Insecticide resistance of vectors | Mean | Phenotypic resistance to insecticides in mosquito species from the An. gambiae complex in Sub-Saharan Africa | Parameterising models of gene drive dynamics in mosquitoes and malaria transmission in humans |

### S2. Variables used in selecting representative environments

To capture variation in prevalence across the region, we used a layer of estimates of the *P. falciparum* prevalence in humans in the year 2019 [1]. We calculated the quartiles of the prevalence values across all grid cells in the region. For each area we then calculated the average malaria prevalence in the human population (Table S2) and set a requirement for the selection of areas that four areas must fall within each of the four prevalence quartiles. In addition, our set of selected areas was chosen to capture variation in mosquito population characteristics. Specifically, we aimed to capture variation in mosquito vector species composition, mosquito abundance and seasonality (see the main text). To represent geographic variation in vector species composition we used fine resolution geospatial layers of predictions of the relative abundance of three vector species groups[2], which we used to estimate species composition for each selected area (Table S2). We used two variables describing rainfall patterns to represent geographic variation in the availability and seasonality of larval habitat. The first variable was long term average annual rainfall across each area[7], and the second variable was the spatial maximum of the average number of dry weeks in a year, over all years in the period 1990-2018 (Table S2). To represent variation in human population size across areas we used data from WorldPop on the number of humans residing in each area[4], using data from the year 2020.

We used hierarchical cluster analysis to group all the 1° by 1° areas within our west African region into a set of four clusters with respect to the above five variables describing mosquito population characteristics. Clusters were identified using hierarchical cluster analysis, implemented using the hclust function from R cluster package, with clustering defined by the average Euclidean distance (using hclust with method= “average”).

### S3. Mosquito metapopulation model

#### S3.1 Overview

In this section we fully describe the assumptions of the mosquito metapopulation model. As stated in the main text, the mosquito population is represented as a metapopulation of sub-populations located at the sites of human settlements, for each species and in each area. Each sub-population is composed of juveniles, adult males, virgin females, and mated females. We assume that larval competition affects larval mortality, and increases with larval density to an extent that depends on the amount of larval habitat in a given sub-population. We assume both adult males and mated females may move between neighbouring populations (*dispersal*).

The first part of this description (section 1.2; *Modelling the landscape*) sets out the assumptions relating to landscape. We first describe how the distribution of settlements in each area (henceforth the *landscape*) is inferred from *World Settlement Footprint* data. We next describe how larval habitat is estimated from data representing rainfall, water courses, the human population spatial distribution and, finally, the relative abundances of each species.

The second part (section 1.3; *Local population dynamics*) describes the dynamics of the mosquito sub-populations and how they are affected by larval habitat. We next describe the assumptions of dispersal between sub-populations. Finally, we describe the assumptions that enable us to consider gene drive dynamics in the model; specifically we set out the assumptions of biased inheritance and fitness effects of a gene drive allele at the doublesex locus.

### S3.2 Modelling the landscape

Here we describe the input data layers that we used to infer settlement locations across the study area, and the variables affecting the spatial and temporal distribution of larval habitat.

##### 3.2.1 Settlement locations

We inferred locations of high human density ("settlements") using World Settlement Footprint (WSF) data ([3]; Table S1). This data demarks the presence or absence of buildings at 10m resolution, and is available for any location globally. To compute settlement locations, we determined the number of built-up 10m by 10m cells within each square kilometre and defined a square kilometre to be a settlement if this number exceeded a threshold of 0.5%. (i.e. each square km is classified as a settlement if the number of built-up sub-cells exceeds 49 of the 10,000 sub-cells within the square). Note that, in our model, the smallest possible distance between neighbouring settlements is 1km, which occurs if neighbouring squares are both identified as settlements.

This method of settlement inference differs from our previous approach which used settlement databases [14]. We opted for this approach because the WSF dataset has been constructed with uniform methods across the region [15].

##### 3.2.2 Settlement localities

After determining the settlement locations in a given region, we partitioned the region into "localities" by applying a Voronoi tessellation (Fig. 1B). As described below, the localities are used to compute the size of the resident human population associated to each settlement, the length of water courses associated to each settlement, and the dispersal between neighbouring settlements.

##### 3.2.3 Human population per settlement

Human distribution data were obtained from "WorldPop" ([4]; Tables S1), which estimates the human population size in 1km by 1km cells across the globe; we used data for the year 2020. For each settlement, we determined the WorldPop cells with centres within the perimeter of its locality, and summed the population estimates of these cells to obtain a human population estimate for the settlement. We denote by $h_{x,a}$ the estimated human population size in location $(x, a)$ where $x$ is the settlement and $a$ is the area. The variable $h_{x,a}$ is a component of estimating mosquito carrying capacity in this settlement, as described below (see equation 1 below).

##### 3.2.4 Water courses

Watercourse data were obtained from "HydroATLAS", which charts rivers and lakes globally [5,6] (Table S1). The data classifies river segments by their average flow rate, and we re-classified all river segments with flow between $0.1$ and $10\ m^3s^{-1}$ as 'seasonal' and segments with flow greater than $10\ m^3s^{-1}$ as 'permanent', though we acknowledge that this classification is approximate. For each settlement $x$ in area $a$, we denote by $W_{1,x,a}$ the total length of seasonal river segments passing through its locality, and by $W_{2,x,a}$ the sum of the total length of permanent river segments and the total length of lake edge in the locality (we assume lake edges are always permanent). Like human population sizes, these variables contribute to calculating the mosquito carrying capacity for each settlement (see equation 1 below).

##### 3.2.5 Rainfall

Rainfall data were obtained from "ERA5" (Table S1 and S2), which estimates hourly rainfall at all points on a spatial lattice with vertices 0.25 degrees ($\approx 27.5km$) apart from historical observations [7]. We aggregated the data to obtain average weekly rainfall within the time period January 1st 1979 to December 31st 2021 at all points on the lattice within our study area (West Africa). We assume the rainfall at a location $(x, a)$ and on day $d$, denoted by $\lambda_{x,a,d}$, is given by the rainfall yearly time-series at the nearest point on the aggregated rainfall data lattice. We assume local carrying capacity increases with rainfall (see equation 1 below).

##### 3.2.6 Species distributions

Mosquito species composition was estimated using a predictive map describing the relative abundance of *An. arabiensis* versus *An. funestus* versus *An. gambiae* and *An. coluzzii* (which are combined) in each ≈ 5 x 5km cell (0.0416 x 0.0416 degrees) across the continent [2]. We mapped this onto the settlement locations, by assigning to each settlement the species abundances of the nearest gridcell of the species distribution layer. We assume the carrying capacity for species $q$ in location $(x, a)$ is proportional to relative abundance, which we denote as $\pi_{x,a,q}$ (see equation 1 below).

### S3.3 Local population dynamics

##### 3.3.1 Juvenile stage

We model all juvenile mosquito stages (egg, larvae, pupae) as a single class called "juveniles" and denote by $J_{x,a,q,d}$ the number of juveniles in location $(x, a)$ and of species $q$ on day $d$. Each day, a juvenile will either die or age by one day until it completes development and becomes an adult.

The time required for each juvenile to develop from egg to adulthood is assigned at random from the range 0 to 19 days with probability $DEV_\tau$ of requiring $\tau$ days. We use as default the distribution given by

$$DEV_{tau} = \begin{cases} 0\ if\ \tau < 10 \\ 0.1\ if\ 10 \leq \tau \leq 19 \end{cases}$$

(1)

such that all juveniles take between 10 and 19 days to develop. Note that fast-developing juveniles are more likely to survive development, so that the adult population will be skewed towards those that developed relatively quickly as juveniles.

The probability that a juvenile of species $q$ in location $(x, a)$ survives day $d$ is

$$p_{x,a,q,d} = \sqrt[15]{\frac{K_{x,a,q,d}}{K_{x,a,q,d} + J_{x,a,q,d}}}(1 - \mu_{JUV}),$$

(2)

where $\mu_{JUV}$ represents density-independent mortality (assumed to be the same for all species groups), and $K_{x,a,q,d}$ is a variable controlling the strength of mortality due to larval competition for resources. The exponent 1/15 is set so a juvenile which requires 15 days to develop will die from larval competition with probability one half, if the number of juveniles in the population is equal to $K_{x,a,q,d}$.

The variable $K_{x,a,q,d}$ varies by location, species, and day, because it is influenced by location-specific factors (the presence of local water bodies, the species distribution and the human population), as well as by rainfall which varies in both time and space. It is computed as:

$$K_{x,a,q,d} = \pi_{x,a,q} DENS^*_{x,a} \left[K_{0,a} + K_{1,a}\left(1 - e^{-\nu_1 \lambda_{x,a,d}}\right) + K_{2,a}\left(1 - e^{-\nu_2\left[W_{1,x,a}\left(1-e^{-\nu_3\lambda_{x,a,d}}\right)+W_{2,x,a}\right]}\right)\right] \tag{3}$$

where $DENS^*_x$ relates to the human population density. We set

$$DENS^*_{x,a} = \begin{cases} DENS_{x,a} = \dfrac{h_{x,a}}{AREA_{x,a}} \text{ if } DENS_{x,a} \le N_H \\ N_H \text{ if } DENS_{x,a} > N_H \end{cases} \tag{4}$$

where $h_{x,a}$ and $DENS_{x,a}$ is the human population size and density at location $(x, a)$ whose area is $AREA_{x,a}$, and $N_H$ is a parameter that ensures mosquito population sizes do not increase indefinitely with human population size. This formulation means that mosquito number increases with the number of humans in small settlements (where the human population density $< N_H$), but will plateau in densely populated settlements such as urban environments.

The three terms within the large square bracket in the equation for $K_{x,a,q,d}$ relate to a baseline level of larval habitat ($K_{0,a}$), the influence of rainfall (scaled by $K_{1,a}$), and the influence of water courses (scaled by $K_{2,a}$), as follows.

We set $K_{0,a} > 0$ to assume there is a baseline amount of larval habitat that is present year-round. This ensures that populations typically remain extant through the dry season, in the absence of gene-drive interventions. In previous work, we have explored alternative explanations for dry season persistence such as aestivation and long distance dispersal [14].

The second term represents the contribution to breeding habitats from rainfall on day $d$, $\lambda_{x,a,d}$, through the creation of puddles and other small water bodies in which anophelines are known to breed. In the absence of rainfall this term is zero, but as precipitation rises it increases at a rate determined by the parameter $\nu_1$ to asymptote at a level $K_{1,a}$. Recall that rainfall is averaged by week so that, for any location $(x, a)$, this contribution to larval habitat varies on a weekly rather than daily basis.

The final term represents the larval sites associated with rivers and lakes which may be seasonal ($W_{1,x,a}$) or perennial ($W_{2,x,a}$). We assume that seasonal water courses are replenished by rainfall at a rate determined by the parameter $\nu_3$, providing the same density of breeding habitat as permanent sites when rainfall is heavy. Mosquito carrying capacity increases at a rate determined by the parameter $\nu_2$ as the density of both types of water body grows, the influence reaching an asymptote at $K_{2,a}$.

##### 3.3.2 Setting the area-specific carrying capacities

For each species group, the maximum daily number of adult female mosquitoes in each area $a$ throughout the entire simulation period, $\hat{F}_{a,q}$, was determined by a model of malaria transmission dynamics (described below) which fitted $\hat{F}_{a,q}$ to match predictions of average annual malaria prevalence in humans in each year to data. For each area $a$, the carrying capacity parameters $K_{1,a}$ and $K_{2,a}$ were adjusted such that population size simulated from the mosquito metapopulation model matched $\hat{F}_{a,q}$. The values of $K_{1,a}$ and $K_{2,a}$ were assumed to be equal for each vector species group.

##### 3.3.3 Adult stage

Juveniles that complete development become male or virgin female adults with equal probability. For a given species $q$ and location $(x, a)$, virgin females mate with probability $\frac{M_{x,a,q,d}}{\beta+M_{x,a,q,d}}$ per day, where $M_{x,a,q,d}$ is the number of males (of that species) in the population and $\beta$ is the adult male population size at which the daily probability of mating is one half. We assume females only mate at most once during their lifetimes. Each mated female lays a Poisson distributed number of eggs per day until she dies, with expectation that depends on her genotype as described below.

Adult males die with a constant probability $\mu_{MALE}$ per day which is assumed to be the same for all species groups. Adult females die with probability $\mu_{FEM_{a,q,d}}$ per day, which may vary by area $a$, species $q$ and day $d$. For each area and species group, a time series $\{\mu_{FEM_{a,q,d}}\}$ over the simulation period was computed based on the annual coverage of ITNs and IRS across the human population residing in each area (see below).

##### 3.3.4 Dispersal

Adult mosquitoes move between neighbouring settlements in two ways, which we will refer to as 'neighbourhood dispersal' and 'border dispersal'. Neighbourhood dispersal represents mosquito movements to settlements within a neighbourhood of radius $L_D$, and occurs at rate $\rho_n$ (the probability that an individual adult mosquito disperses on a given day). We did not consider species-specific differences in this parameter, though for all species groups we simulated the model across a wide range of values to account for uncertainty in dispersal propensity (see Results). If an individual at location $(x_1, a)$ does disperse in this way, the probability that it moves to location $(x_2, a)$ is

$$\frac{L_D - DIST_{x_1,x_2,a}}{\sum_k (L_D - DIST_{x_1 x_k,a})}, \tag{5}$$

where $DIST_{x_1 x_2,a}$ is the distance between the two settlements and the sum is over all settlements within the $L_D$ radius (not including the focal settlement). This formulation means that individuals move more frequently between nearby than distant sites.

Border dispersal represents additional movements of mosquitoes between settlements whose localities share a border, and occurs at rate $\rho_b$ (as with neighbourhood dispersal, we do not consider species-specific differences in this parameter). The selection of a destination from these dispersal events is based on the length of shared border. If the initial location is $x_1$, the probability that destination $x_2$ is selected is

$$LEN_{x_1 x_2,a} \Big/ \sum_k LEN_{x_1 x_k,a} \tag{6}$$

where $LEN_{x_1 x_2}$ is the length of the locality border shared by focal settlement $x_1$ and settlement $x_2$ and the sum is over all settlements sharing a border with the focal settlement. The inclusion of locality border dispersal in our model ensures that no settlement is entirely unconnected.

##### 3.3.5 Gene drive

Following North et al [16], the model considers three types of allele at the doublesex locus: wildtype (W), gene-drive (G), and a non-functional mutant allele (r2 allele; X) that is resistant to homing. There are therefore six possible mosquito genotypes (W/W, W/G, G/G, W/X, X/X, G/X) and, for each species group, the model keeps track of the numbers of mosquitoes with each genotype in each population. The model also tracks the genotype of the mate of mated females to determine the genotype frequencies in her offspring, resulting in 36 potential types of mated adult female. For each species group, homing occurs during meiosis at a rate $\boldsymbol{e}$, leading to gene drive alleles representing $\frac{\mathbf{1+e}}{\mathbf{2}}$ of the gametes produced by heterozygous (W/D) males and females. Of the remaining $\frac{\mathbf{1-e}}{\mathbf{2}}$ fraction of gametes, a fraction $1 - \chi$ are wildtype, and $\chi$ are converted into X alleles by NHEJ repair.

Females that are either fully wildtype (W/W) or heterozygous for the resistant allele (W/X) are assumed to have maximal fertility of $l$ (expected number of eggs per day) owing to having at least one functional doublesex gene (we assume this is the same for each species group). The relative fitness of females that are heterozygous for the gene drive (W/G) is $\omega$ ($0 \leq \omega \leq 1$) such that their fertility is $\omega \times l$. The reduced fertility of these individuals may be caused by expression of the transgene in somatic cells, or by parental deposition of the cas9 endonuclease in the sperm of egg [17,18]. Females that lack a copy of the wildtype allele (G/G, X/X, and G/X) are assumed to be sterile. We assume there are no other effects of genotype.

##### 3.3.6 Parameters of models of mosquito and gene drive dynamics

The parameters of the mosquito meta-population model representing the spread dynamics of population suppression gene drives are provided in Table S3.

**Table S3.** Parameters, variables, and mathematical notation used in the mosquito population model

| Parameter/variable | Symbol | Estimate | Units | Reference |
|---|---|---|---|---|
| **General indices** | | | | |
| Locality | $x$ | n/a | n/a | n/a |
| Day | $d$ | n/a | n/a | n/a |
| Year | $y$ | n/a | n/a | n/a |
| Species | $q$ | n/a | n/a | n/a |
| Area | $a$ | n/a | n/a | n/a |
| Larval duration | $\tau$ | n/a | n/a | n/a |
| **Genotypes** | | | | |
| Wildtype type mosquitoes | $W$ | n/a | n/a | n/a |
| Gene drive mosquitoes | $G$ | n/a | n/a | n/a |
| Resistant mosquitoes | $X$ | n/a | n/a | n/a |
| **Gene drive parameters** | | | | |

| | | | | |
|---|---|---|---|---|
| Homing rate | $e$ | 0.95 | $generation^{-1}$ | [18] |
| Proportion of non-homed alleles that form non-functional resistant alleles (R2 alleles) | $\chi$ | 0.5 | n/a | [17] |
| Relative fitness | $\omega$ | variable | n/a | [17] |
| **Life-history parameters** | | | | |
| Larval density independent mortality | $\mu_{JUV}$ | 0.05 | $day^{-1}$ | [14] |
| Probability that juvenile requires $\tau$ days to develop | $DEV_\tau$ | $\begin{cases} 0\ if\ \tau < 11 \\ 0.1\ if\ 11 \le \tau \le 20 \end{cases}$ | n/a | [19,20] |
| Male mosquito mortality | $\mu_{MALE}$ | 0.125 | $day^{-1}$ | [14] |
| Female mosquito mortality | $\mu_{FEM_{a,q,d}}$ | variable | $day^{-1}$ | see Methods |
| Oviposition rate | $l$ | 9 | $day^{-1}$ | [14] |
| **Carrying capacity parameters** | | | | |
| Baseline contribution to larval habitat | $K_{0,a}$ | 20 | n/a | [14] |
| Maximum contribution from rain *per se* | $K_{1,a}$ | variable | n/a | see Methods |
| Maximum contribution from water bodies | $K_{2,a}$ | variable | n/a | see Methods |
| Increase in $K_0/K_1$ per mm rain per week (when rainfall low) | $\nu_1$ | 0.03 | $(mm\ rain)^{-1}(week)^{-1}$ | [14] |
| Increase in $K_0/K_2$ per km standing water (within locality; when water bodies rare) | $\nu_2$ | 0.8 | $(km\ water)^{-1}$ | [14] |
| Increase in length of standing water from non-permanent waterways per km non-permanent waterways (within locality) per mm rain | $\nu_3$ | 0.03 | $(mm\ rain)^{-1}(week)^{-1}$ | [14] |

| | | | | |
|---|---|---|---|---|
| per week (when rainfall low) | | | | |
| Threshold human density | $N_H$ | 100 | $(people)(km)^{-2}$ | Expert judgement |
| Number of males when females mate with probability ½ per day | $\beta$ | 100 | n/a | [14] |
| **Dispersal parameters** | | | | |
| Dispersal neighbourhood radius | $L_D$ | 10 | km | Expert judgement |
| Local dispersal probability | $\rho_n$ | 0.001-0.025 | $(day)^{-1}$ | [14] |
| Border dispersal probability | $\rho_b$ | $\rho_n/100$ | $(day)^{-1}$ | Expert judgement |
| **Variables** | | | | |
| Settlement area | $AREA_{x,a}$ | n/a | $(km)^{-2}$ | n/a |
| Human population in settlement | $h_{x,a}$ | n/a | $(people)$ | n/a |
| Human density in settlement | $DENS_{x,a}$ | n/a | $(people)(km)^{-2}$ | n/a |
| Adjusted human density in settlement | $DENS^*_{x,a}$ | n/a | n/a | n/a |
| River segments seasonal | $W_{1,x,a}$ | n/a | $(km)$ | n/a |
| River segments permanent | $W_{2,x,a}$ | n/a | $(km)$ | n/a |
| Rainfall | $\lambda_{x,a,d}$ | n/a | $(mm)(week)^{-2}$ | n/a |
| No. female adult mosquitoes | $F_{x,a,q,d}$ | n/a | $(mosquitoes)$ | n/a |
| No. male adult mosquitoes | $M_{x,a,q,d}$ | n/a | $(mosquitoes)$ | n/a |
| No. juveniles | $J_{x,a,q,d}$ | n/a | $(mosquitoes)$ | n/a |
| Larval survival | $p_{x,a,q,d}$ | n/a | $(day)^{-1}$ | n/a |
| Fraction of species | $\pi_{x,a,q}$ | n/a | n/a | [2] |
| Distance between two settlements | $DIST_{x_1,x_2,a}$ | n/a | $km$ | n/a |
| Length of border shared between two settlements | $LEN_{x_1,x_2,a}$ | n/a | $km$ | n/a |
| Maximum daily number of adult female mosquitoes in area | $\hat{F}_{a,q}$ | n/a | $(mosquitoes)$ | n/a |

### S4. Modelling malaria infection dynamics in humans

Our model of malaria infection dynamics in humans assumes that the human population within each modelled area $a$ is well-mixed, such that spatial variation in individual infection risk is not considered, and closed, such that immigration and emigration of infections to and from the area is not considered. We model malaria infection and immunity dynamics in humans using the 'malariasimulation' model [21], which is an open-source R-package that implements an individual-based model of malaria infections in humans [22-25]. The full mathematical formulation of 'malariasimulation' is published elsewhere [25,26] and here we provide a summary of the key details. The parameter values that we use in 'malariasimulation' to model human infection and immunity are given in Table S4.

**Table S4.** Baseline parameters used in 'malariasimulation' to model malaria infection and immunity in humans.

| Parameter | Symbol | Estimate | Reference |
|---|---|---|---|
| Human infection duration | | | |
| Latent period | $d_l$ | 12 days | [27] |
| Patent infection | $d_A$ | 200 days | [27,28] |
| Clinical disease (treated) | $d_T$ | 5 days | [29] |
| Clinical disease (untreated) | $d_D$ | 5 days | [30] |
| Sub-patent infection | $d_U$ | 110 days | [25] |
| Immunity decay rates | | | |
| Inverse of decay rate for maternal immunity to clinical disease | $d_M$ | 67.6952 days | [22] |
| Inverse of decay rate for maternal immunity to severe disease | $d_{VM}$ | 76.8365 days | [24] |
| Inverse of decay rate for acquired pre-erythrocytic immunity | $d_B$ | 10 years | [23] |
| Inverse of decay rate for acquired immunity to clinical disease | $d_{cA}$ | 30 years | [31] |
| Inverse of decay rate for acquired immunity to severe disease | $d_{vA}$ | 30 years | [31] |
| Inverse of decay rate for acquired immunity to detectability | $d_{ID}$ | 10 years | [23] |
| Maternal immunity | | | |
| New-born clinical immunity relative to mother | $I_{CM}$ | 0.774368 | [22] |
| New-born severe immunity relative to mother | $I_{VM}$ | 0.195768 | [24] |
| Probability of pre-erythrocytic infection | | | |
| Maximum probability due to no immunity | $b_0$ | 0.59 | [22] |
| Maximum reduction due to immunity | $b_1$ | 0.5 | [23] |
| Scale parameter | $I_{B0}$ | 43.9 | [22] |
| Shape parameter | $\kappa_B$ | 2.16 | [22] |
| Probability of clinical infection | | | |

| | | | |
|---|---|---|---|
| Maximum probability due to no immunity | $\phi_0$ | 0.792 | [22] |
| Maximum reduction due to immunity | $\phi_1$ | 0.00074 | [22] |
| Scale parameter | $I_{C0}$ | 18.02366 | [22] |
| Shape parameter | $\kappa_C$ | 2.36949 | [22] |
| Probability of severe infection | | | |
| Maximum probability due to no immunity | $\theta_0$ | 0.074989 | [24] |
| Maximum reduction due to immunity | $\theta_1$ | 0.000119 | [24] |
| Scale parameter | $I_{V0}$ | 1.09629 | [24] |
| Shape parameter | $\kappa_V$ | 2.00048 | [24] |
| Age dependent modifier | $f_{V0}$ | 0.141195 | [24] |
| Age dependent modifier | $\alpha_V$ | 2493.41 | [24] |
| Age dependent modifier | $\gamma_V$ | 2.91282 | [24] |
| Immunity reducing probability of detection | | | |
| Time-scale at which immunity changes with age | $f_{D0}$ | 0.007055 | [22] |
| Scale parameter relating age to immunity | $g_D$ | 21.9 years | [22] |
| Shape parameter relating age to immunity | $\gamma_D$ | 4.8183 | [22] |
| Minimum probability of detection | $d_{min}$ | 0.160527 | [22] |
| Scale parameter | $I_{D0}$ | 1.577533 | [22] |
| Shape parameter | $\kappa_D$ | 0.476614 | [22] |
| Periods over which immunity is not boosted | | | |
| Period in which pre-erythrocytic immunity cannot be boosted | $u_B$ | 7.2 days | [22] |
| Period in which clinical immunity cannot be boosted | $u_C$ | 6.06 days | [22] |
| Period in which severe immunity cannot be boosted | $u_V$ | 11.4321 days | [24] |
| Period in which immunity to detectability cannot be boosted | $u_D$ | 9.44512 days | [22] |
| Infectivity towards mosquitoes | | | |
| Infectivity of clinically diseased humans | $c_D$ | 0.068 | [32] |
| Infectivity of humans with sub-patent infection | $c_U$ | 0.0062 | [25] |
| Infectivity of treated humans | $c_T$ | 0.021896 | [33] |
| Age and heterogeneity | | | |
| Age dependent biting parameter | $g_0$ | 8 years | [34,35] |
| Age dependent biting parameter | $\rho$ | 0.85 | [34,35] |
| Variance of the log heterogeneity in biting rates | $\sigma^2$ | 1.67 | [36] |

| Human demographic and behavioural parameters | | | |
|---|---|---|---|
| The average age of humans | | 22 years | [37] |
| Probability of seeking treatment | $f_{T,a}^{y}$ | Varies by area *a* and year *y* | Tables S1, S2 |
| *Plasmodium* infection dynamics in mosquitoes | | | |
| Extrinsic incubation period | $T_E$ | 10 days | [38] |
| Daily rate of seeking a blood meal | $f$ | 0.33 | [39] |
| Proportion of bloodmeals taken on humans | $Q_0$ | 0.92 (*An. gambiae)*<br>0.71 (*An. arabiensis)*<br>0.94 (*An. funestus*) | [40] (within the maximum and minimum values reported) |

Susceptible individuals ($i$) in area *a* in year *y* experience a force of infection $\Lambda_{i,a}^{y}$ which depends on their exposure to bites from infectious mosquitoes as well as their level of pre-erythrocytic immunity. New-born individuals have a level of maternally-acquired immunity that decays over the first six months of life. Following infection, individuals experience a delay of length $d_I$ representing the period from infection to appearance of blood-stage infection, after which they may either develop clinical disease or asymptomatic infection (*A*). Individuals who develop clinical disease be treated with a probability $f_{T,a}^{y}$, moving to infection state *T*, or will not seek treatment (with probability $1 - f_{T,a}^{y}$) moving to infection state *D*. Those in the latter state eventually resolve symptoms and move to the asymptomatic state (A). Those that are treated subsequently experience a period of drug-dependent partial protection from re-infection (modelled as a Weibull survivorship function) before returning to the fully susceptible state (S). From the asymptomatic state, as parasite density gradually reduces due to the immune response, asymptomatic individuals move to the sub-patent infection state (U) after which they clear infection and return to the susceptible state S. Individuals in states D, A and U can be re-infected (super-infection) and will move into infection states D, T or A following the same process as for primary infection. The transition pathways between these states are depicted in Figure S1.

We use national life-tables to model death from natural causes [37]. Individuals have age-specific rates at which that are removed from the population that match national life-table rates. Deaths associated with malaria are tracked separately [41]. A new-born individual replaces an individual who has died with matched heterogeneity levels for exposure to infection. In this manner the population size in the simulation remains constant.

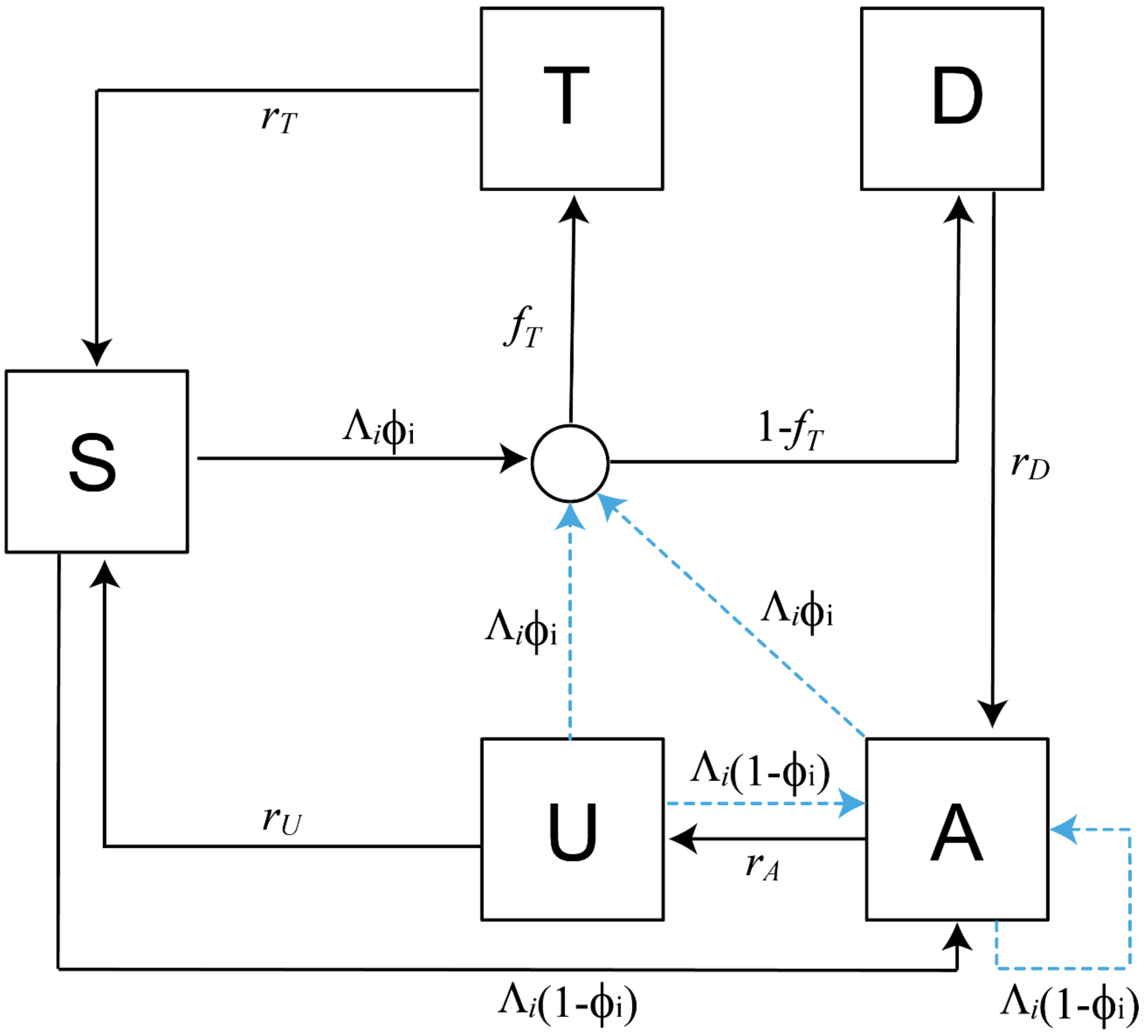

**Figure S1. Progression between human infection states**. States are shown in boxes and state transitions by arrows with associated hazard rates. The circle represents the treatment node. Superinfection is indicated by the dashed blue arrows. S = susceptible, D = clinical disease, T = successfully treated disease, A = asymptomatic patent infection, U = asymptomatic sub-patent infection. Diagram is reproduced from SI Appendix to Winskill et al. [26].

*Heterogeneity in human biting rates*

Individual humans experience a unique biting rate which depends on the product of their age-dependent biting rate, $\psi(g)$ and their relative biting rate $\zeta_i$ . The relative biting rate for age *g* is defined as:

$$\psi(g) = 1 - \rho\exp\left(-\frac{g}{g_0}\right) \tag{7}$$

where $\rho$ and $g_0$ are parameters that determine how biting rates increase with age due to increases in body size.

The relative biting rate is drawn from a Log-normal distribution with a mean of 1:

$$\log(\zeta_i) \sim N\left(\frac{-\sigma^2}{2}, \sigma^2\right)$$

( 8 )

An individual *i* with an age $g$ at time $t$ then experiences an EIR $\varepsilon_i(g,t)$ of

$$\varepsilon_i(g,t) = \varepsilon_0(g,t)\zeta_i\psi(g)$$

( 9 )

where $\varepsilon_0(t)$ is the mean EIR experienced by adults at time *t*.

*Immunity*

Human immunity to infection is modelled as a dynamic process that depends on age and exposure to infection. The maternal immunity that is acquired by infants provides some protection from clinical disease and severe disease at birth is denoted $I_{CM}$ and $I_{VM}$ respectively. These forms of protective maternal immunity decays exponentially at constant rates $r_M = 1/d_M$ and $r_{VM} = 1/d_{VM}$, respectively.

Acquired immunity to infection (pre-erythrocytic immunity) develops at older ages, is boosted by one level following each infected bite provided it is at least $u_B$ days since the last exposure, and decays exponentially in between exposures with rate $r_B = 1\backslash d_B$. Effectively, acquired immunity protects a person from developing patent infection following an infectious bite.

Blood stage immunity is assumed to control parasite density and hence affect the probability of developing severe disease, clinical disease and the detectability of asymptomatic infection. Acquired immunity to each of severe disease, clinical disease and detectability of infection is tracked separately, is boosted by one level following each patent infection provided it is at least $u_V$, $u_C$, $u_D$ days respectively since the last exposure, and decays exponentially between exposures with rate $r_{vA} = 1\backslash d_{VA}$, $r_{cA} = 1\backslash d_{cA}$ and $r_{ID} = 1\backslash d_{ID}$ respectively.

All immunity levels are converted to individual time-dependent probabilities using Hill functions. The probability that individual i who is exposed to an infectious bite at time t develops a patent infection is given by

$$b_i(t) = b_0\left(b_1 + \frac{1-b_1}{1+\left(\frac{I_B(i,t)}{I_{B_0}}\right)^{\kappa_B}}\right)$$

( 10 )

where $b_0$ is the probability of infection with no immunity, $b_0\ b_1$ is the minimum probability, $I_{B0}$ and $\kappa_B$ are scale and shape parameters respectively and $I_B(i,t)$ is the level of pre-erythrocytic immunity of individual *i* at time *t.*

The probability that an individual *i* who has been infected develops clinical disease at time *t* is

$$\phi(t) = \phi_0\left(\phi_1 + \frac{1-\phi_1}{1+\left(\frac{I_{CA}(i,t)+I_{CM}(i,t)}{I_{C0}}\right)^{\kappa_c}}\right)$$

( 11 )

where $\phi_0$ is the probability of disease with no immunity and $\phi_0\phi_1$ is the minimum probability, and $I_{c0}$ and $\kappa_c$ are scale and shape parameters respectively. Acquired immunity to clinical disease is $I_{CA}(i,t)$ and $I_{CM}(i,t)$ respectively.

The probability that an individual *i* who has been infected develops severe disease at time *t* is

$$\theta_i(g,t) = \theta_0\left(\theta_1 + \frac{1-\theta_1}{1 + f_v(i,g)\left(\frac{I_{VA}(i,t) + I_{VM}(i,t)}{I_{V0}}\right)^{\kappa_v}}\right)$$

( 12 )

where $\theta_0$ is the probability of disease with no immunity and $\theta_0\theta_1$ is the minimum probability, and $I_{V0}$ and $\kappa_v$ are scale and shape parameters respectively. $I_{VA}(i,t)$ is the level of acquired immunity to severe disease and $I_{VM}(i,t)$ is the level of maternal immunity to sever disease of individual *i* at time *t* and

$$f_v(i,g) = 1 - \frac{1 - f_{V0}}{1 + \frac{\alpha}{\alpha_V}^{\gamma_V}}$$

( 13 )

is an age-dependent modifier of the risk of severe disease, where $f_{V0}$, $\alpha_V$ and $\gamma_V$ are parameters.

Parasite infection can be detected by microscopy. There is a minimum probability of detection $d_{min}$ of an asymptomatic infection in individual *i* of age *g* at time *t* . The capacity to detect infection is determined by:

$$q_i(g,t) = d_1 + \frac{1 - d_{min}}{\left(\frac{1 + I_D(i,t)}{I_{D0}}\right)^{\kappa_D} f_D(i,g)}$$

( 14 )

$I_{D0}$ and $\kappa_D$ are scale and shape parameters respectively, $I_D(i,t)$ is the level of acquired immunity to the detectability of infection of individual *i* at time *t* and

$$f_D(i,g) = 1 - \frac{1 - f_{D0}}{1 + \left(\frac{g}{g_D}\right)^{\gamma_D}}$$

( 15 )

represents an age-dependent modifier for infection detectability, where $f_{D0}$, $g_D$ and $\gamma_D$ are constant parameters.

*Infectivity towards mosquitoes*

Individuals in different infection states have a different level of infectiousness towards mosquitoes, with the highest infectivity associated with the states in which parasite density is highest (i.e. disease). Onwards infectiousness is $c_A$, $c_D$ and $c_U$ in states $A$, $D$and $U$ respectively, and $c_T$ following treatment.

*Stochastic equations*

The model is implemented as a stochastic individual-based model using Komolgorov forward equations. The full model equations are provided in [40] following [25] and are reproduced here for completeness. Any individual *i* at time *t* can be placed into a state $\{j, t_T, k, t_k, l, t_l, m, t_m, n, t_n, g, t\}$, where their age $g$, and infection status *j* (*S, D, A, U, T*), are tracked across time. Here, $t_T$ represents the time of last treatment for individual *i*, *k* depicts the level of infection-blocking immunity and $t_k$ is the time at which infection blocking immunity was last boosted (relating to the last time this individual was exposed to an infectious mosquito bite). The level of the most recent boost of naturally-acquired immunity is denoted $l$ and the time when the exposure occurred $t_l$ . The level of parasite detection immunity is denoted $m$ and the time is $t_m$ . Likewise, *n* and $t_n$ represent the boost and time of exposure for severe disease immunity. The notation $I_{mat}$ and $n_{mat}$ represent these levels and time for maternally-acquired immunity to clinical disease and severe diseases respectively. The Kronecker delta $\delta_{p,q} = 1\ if\ p = q$ and 0 otherwise.

We let the probability density function be denoted as $P_i(j, t_T, k, t_k, I_{mat}, l, t_l, m, t_m, n_{mat}, n, t_n, g, t)$ for individual i in state $\{j, t_T, k, t_k, I_{mat}, l, t_l, m, t_m, n_{mat}, n, t_n, g, t\}$ at time *t* . Then, the time evolution of the system can be written by the following forward equation:

**1** $$\frac{\partial P_i(j, t_T, k, t_k, I_{mat}, l, t_l, m, t_m, n_{mat}, n, t_n, g, t)}{\partial t} + \frac{\partial P_i(j, t_T, k, t_k, I_{mat}, l, t_l, m, t_m, n_{mat}, n, t_n, g, t)}{\partial a} =$$

**2** $$\delta_{j,S}[r_t P_i \ + r_U P_i(U, t_T, k, t_k, I_{mat}, l, t_l, m, t_m, n_{mat}, n, t_n, g, t)] +$$

**3** $$\delta_{j,A}\left[r_D\left(P_i(D, t_T, k, t_k, I_{mat}, l, t_l, m, t_m, n_{mat}, n, t_n, g, t)\right)\right] +$$

**4** $$\delta_{j,U}\left[r_D\left(P_i(A, t_T, k, t_k, I_{mat}, l, t_l, m, t_m, n_{mat}, n, t_n, g, t)\right)\right] +$$

**5** $$(1 - b_i)\varepsilon_i(t - d_l)\left[\delta_{j,S} + \ \delta_{j,D} + \delta_{j,A} + \delta_{j,U}\right]\vartheta_b \circ P_i(j, t_T, k, t_k, I_{mat}, l, t_l, m, t_m, n_{mat}, n, t_n, g, t) +$$

**6** $$b_i\varepsilon_i(t - d_l)\left[\delta_{j,A}\left(1 - \phi_i(t)\right) + \delta_{j,D}\phi_i(t)(1 - f_T) + \delta_{j,T}\phi_i(t)f_T\right] \vartheta_I \circ \vartheta_b \circ \vartheta_c \circ \vartheta_d \circ \vartheta_v \circ \sum_{j'} s_{i,j'}(t - t_T)P_i(j', t_T, k, t_k, I_{mat}, l, t_l, m, t_m, n_{mat}, n, t_n, g, t) +$$

**7** $$\left[r_B k\frac{\partial}{\partial k} + r_{CA} I\frac{\partial}{\partial I} + r_M I_{mat}\frac{\partial}{\partial I_{mat}} + r_{ID} m\frac{\partial}{\partial m} + r_M n_{mat}\frac{\partial}{\partial n_{mat}} + \ r_{VA} n\frac{\partial}{\partial n}\right] P_i(j, t_T, k, t_k, I_{mat}, l, t_l, m, t_m, n_{mat}, n, t_n, g, t) +$$

**8** $$\mu(g)\delta(g)\delta\left(t_t + T_{big}\right)\delta\left(t_k + T_{big}\right)\delta\left(t_l + T_{big}\right)\delta\left(t_m + T_{big}\right)\delta\left(t_n + T_{big}\right)\delta\left(I_{mat} - mat_c(\zeta_i)\right)\left(n_{mat} - mat_v(\zeta_i)\right)\ \delta_{i,S}\delta_{k,0}\delta_{l,0}\delta_{m,0}\delta_{n,0}P_i(j, t_T, k, t_k, I_{mat}, l, t_l, m, t_m, n_{mat}, n, t_n, g, t) -$$

**9** $$\left[\mu(g) + r_U S_{j,U} \ + r_D S_{j,D} \ + r_A S_{j,A} \ + r_T S_{j,T} \ + h_i(t - t_E)s_{i,j'}(t - t_T)\right]P_i(j, t_T, k, t_k, I_{mat}, l, t_l, m, t_m, n_{mat}, n, t_n, g, t)$$

*( 16 )*

The pathways in Figure S1 are represented in lines 2-4; the flows into the human compartments (T, U, D and A). The boosting of pre-erythrocytic immunity following exposure is shown in line 5 (for the situation where exposure does not lead to an infection). Line 6 captures the exposure leading to infection of people in states S, A or U (with onward movement into states A, D or T). The density of individuals in each of the states of immunity are tracked by the communitive integral operators. These operators $\vartheta_I$, $\vartheta_b$ , $\vartheta_c$, $\vartheta_d$and $\vartheta_v$ are defined below. Line 7 represents the deterministic exponential decay of the respective immunity types. Birth and death processes are indicated in lines 8 and 9.

The commutative integral operators have the following effect on density

$f(j, t_T, k, t_k, I_{mat}, l, t_l, m, t_m, n_{mat}, n, t_n, g, t)$:

$$\vartheta_b \circ f = \delta(t - t_k) \int_0^\infty f(j, t_T, k-1, t_k - u_b - \tau, I_{mat}, l, t_l, m, t_m, n_{mat}, n, t_n, g, t) d\tau + \Phi\left(\frac{t - t_k}{u_b}\right) f(j, t_T, k, t_k, I_{mat}, l, t_l, m, t_m, n_{mat}, n, t_n, g, t)$$

$$\vartheta_c \circ f = \delta(t - t_k) \int_0^\infty f(j, t_T, k, t_k, I_{mat}, l-1, t_l - u_c - \tau, m, t_m, n_{mat}, n, t_n, g, t) d\tau + \Phi\left(\frac{t - t_l}{u_c}\right) f(j, t_T, k, t_k, I_{mat}, l, t_l, m, t_m, n_{mat}, n, t_n, g, t)$$

$$\vartheta_d \circ f = \delta(t - t_k) \int_0^\infty f(j, t_T, k, t_k, I_{mat}, l, t_l, m-1, t_m - u_d - \tau, n_{mat}, n, t_n, g, t) d\tau + \Phi\left(\frac{t - t_d}{u_d}\right) f(j, t_T, k, t_k, I_{mat}, l, t_l, m, t_m, n_{mat}, n, t_n, g, t)$$

$$\vartheta_v \circ f = \delta(t - t_k) \int_0^\infty f(j, t_T, k, t_k, I_{mat}, l, t_l, m, t_m, n_{mat}, n-1, t_n - u_v - \tau, g, t) d\tau + \Phi\left(\frac{t - t_n}{u_v}\right) f(j, t_T, k, t_k, I_{mat}, l, t_l, m, t_m, n_{mat}, n, t_n, g, t)$$

(17)

In these actions, $\Phi(x)$ is an indicator function where

$$\Phi(x) = \begin{matrix} 1 & x < 1 \\ 0 & \text{otherwise} \end{matrix}$$

At birth, *k, l, m* and *n* are set to zero for each individual while $t_l, t_T, t_k, t_m$ and $t_n$ are set to a large negative value $-T_{big}$ (this indicates that there has been no exposure or infection yet). If the previous boost to *k, l, m* and *n* occurred longer than $u_b, u_c, u_d$ and $u_v$ days ago, the naturally-acquired immunity term increase by 1 for each individual receiving an infectious bite (*k*) or who is infected (*l,m* and *n*) respectively. The levels of immunity in individuals aged between 15 and 35 years and within the matched heterogeneity group $\zeta_i$ (termed $mat_c(\zeta_i)$ and $t_v(\zeta_i)$ ) for clinical and severe disease immunity respectively) are used to draw the maternal immunity levels $I_{mat}$ and $n_{mat}$ which are not boosted by exposure. In all cases, immunity levels decay exponentially in the model.

Susceptibility to infection depends on the infection-state and this tracks the prophylactic period of the last drug received. This is defined as:

$$s_{i,j'}(t - t_T) = \begin{matrix} 1 & j' = D, A, U \\ W(t - t_T) & j' = S \\ 0 & j' = T \end{matrix}$$

Here, $W(t-t_T)$ is a Weibull survivorship function modelled to decay over time. The function is specific to the ACT treatment administered.

### S4.1 Malaria infection dynamics in mosquitoes

Our model of mosquito population dynamics is a modification of the 'malariasimulation' model (version 1.4.3) such that the rate of emergence of adult mosquitoes is calculated using our spatial metapopulation model and passed to 'malariasimulation' as an external variable, as described in the Methods section of the main text. Here we provide the mathematical formulation of our mosquito population model.

The total number of adult female mosquitoes of species *q* in area *a* at time *t*, $F_a^q(t)$, is subdivided into the numbers that are not infected with the *P. falciparum* parasite, $F_{S,a}^q(t)$, infected with parasites that are still incubating and not yet transmissible, $F_{E,a}^q(t)$, and infectious with transmissible parasites, $F_{I,a}^q(t)$. The dynamics of these three groups of mosquitoes are given by:

$$\frac{dF_{S,a}^q(t)}{dt} = 0.5e_{a,q}(t) - \mu_{FEM_{a,q}}(t)F_{S,a}^q(t) - \Lambda_a^q(t)F_{S,a}^q(t)$$

$$\frac{dF_{E,a}^q(t)}{dt} = \Lambda_a^q(t)F_{S,a}^q(t) - e^{-\mu_{FEM_{a,q}}T_E}F_{E,a}^q(t)(t-T_E) - \mu_{FEM_{a,q}}(t)F_{E,a}^q(t)$$

$$\frac{dF_{I,a}^q(t)}{dt} = e^{-\mu_{FEM_{a,q}}T_E}F_{E,a}^q(t)(t-T_E) - \mu_{FEM_{a,q}}(t)F_{I,a}^q(t)$$

( 18 )

where $e_{a,q}(t)$ is the daily rate of emergence of adult mosquitoes (see the Methods section in the main text), $\mu_{FEM_{a,q}}(t)$ is the daily rate of adult female mosquito mortality (Table S3), $\Lambda_a^q(t)$ is the daily rate at which species *q* becomes infected with *P. falciparum* in area *a,* and $T_E$ is the entomological inoculation period of the *P. falciparum* parasite (Table S4).

The rate at which adult female mosquitoes become infected is a function of the infectiousness of the human population including an appropriate time-lag $d_l$ to account for the period between humans becoming infected and becoming infectious. The force of infection towards mosquitoes is therefore dependent on all human infected states and is defined as:

$$\Lambda_a^q(t) = \frac{\alpha}{\omega}\iint_0^\infty \zeta\psi(g)(c_D D(\zeta,g,t-d_l) + c_T T(\zeta,g,t-d_l) + c_A A(\zeta,g,t-d_l) + c_U U(\zeta,g,t-d_l))dgd\zeta$$

( 19 )

where $\alpha$ is the biting rate on humans

$$\alpha = Q_0 f$$

$Q_0$ quantifies the level of anthropophagy and *f* is the daily rate of taking blood-meals (see Table S4). The parameter $\omega$ represents a normalising constant for the biting rate over all ages:

$$\omega = \int_0^\infty \psi(g)G(g)dg$$

( 20 )

where *G(g)* is the human age distribution.

### S4.2 Estimating vector abundances

We estimate $e_a$, the maximum daily number of adult females that emerge in area *a,* to match model predictions of the annual average prevalence of *P. falciparum* malaria in children aged between 2-10 years (inclusive) for each year in the pre-gene drive period to the yearly data on prevalence [1] (see the main text). Here we give details of the maximum likelihood method used to estimate $\mathrm{e_a}$.
The prevalence for area a in year y obtained from the geospatial layers, $p^y_{MAP,a}$, is calculated as

$$p^y_{MAP,a} = \frac{1}{H^y_a}\sum_{p=1}^{P} p^y_{MAP,p} H^y_{a,p}$$

( 21)

where $p^y_{MAP,p}$ is the prevalence in pixel *p* in year *k,* $H^y_{a,p}$ is the number of humans residing in pixel *p* in year *y*, and *P* is the total number of pixels in each area. The area-wide number of humans residing in each area in year *y is* denoted $H^y_a$.

We fit the model to maximise a modified likelihood function *L* in which more recent observations are more highly weighted compared to earlier observations[42]:

$$L(\{I^y_a\}|\{p^y_{MAP,a}\}) = \prod_{y=0}^{T-1}\left[Bin\left(H^{T-y}_a, p^{T-y}_{MAP,a}\right)\right]^{\gamma^y}, \quad \alpha \in [0,1]$$

( 22)

where $I^y_a$ is the number of individuals containing blood stage malaria parasites in the human population residing in area *a* in year *y,* and *T* is the final year of the pre-gene drive period. The parameter $\gamma$ upweights more recent observations, and was set to a value of $\gamma$=0.2. This value of $\gamma$ was found to produce the minimum out-of-sample root mean square error (RMSE) between the model-predicted malaria prevalences and the data values $p^y_{MAP,a}$ across all areas *a* in the most recent year of the pre-gene drive period, *T.*

### S4.3 Treatment and Seasonal Malaria Chemoprevention

As described in the main text, the proportion of clinical cases in each area who receive treatment in a given year, $f^y_{T,a}$, receive either artemether-lumefantrine (AL; a form of artemisinin combination therapy (ACT)) or sulphadoxine-pyrimethamine and amodiaquine (SP-AQ). The probabilities of receiving each treatment type are $ACT^k_a$ and $SPAQ^k_a$ (Tables S1 & S2). In each area and year, a proportion of children aged 2-5 years receive seasonal malaria chemoprevention (SMC), dentoed $SMC^y_a$(Table 1) with the drug SP-AQ. We use the 'set_clinical_treatment' function in 'malariasimulation' to implement the pharmacokinetic-pharmacodynamic (PKPD) models of the time-varying efficacy of each type of drug developed by Okell et al[43], using the parameter values in Table S5.

**Table S5**. Parameters of the pharmacokinetic-pharmacodynamic model for the time-varying efficacy of artemether–lumefantrine (AL) and sulphadoxine-pyrimethamine and amodiaquine (SP-AQ).

| Parameter | Definition | Value | Source |
|---|---|---|---|
| $P_{min,AL}$ | Infection probability at infinitely high blood AL concentration | 0.95 | [43] |
| $c_{T,AL}$ | Infectiousness after AL treatment relative to an untreated infection | 0.05094 | [43] |
| $w_{AL}$ | Shape parameter for AL PKPD model | 11.3 | [43] |
| $\lambda_{AL}$ | Scale parameter for AL PKPD model | 10.6 | [43] |
| $P_{min,SPAQ}$ | Infection probability at infinitely high blood SPAQ concentration | 0.9 | [43] |
| $c_{T,SPAQ}$ | Infectiousness after SPAQ treatment relative to an untreated infection | 0.32 | [43] |
| $w_{SPAQ}$ | Shape parameter for SPAQ PKPD model | 4.3 | [43] |
| $\lambda_{SPAQ}$ | Scale parameter for SPAQ PKPD model | 38.1 | [43] |

### S4.4 Insecticide-treated bednets

#### 4.4.1 Impacts on vector mortality and blood-feeding

Our model of the impacts of ITNs on rates of mortality and human blood feeding in the vector population is implemented using the 'set_bednets' function in 'malariasimulation' and is provided on [21]. This model considers that two possible types of ITNs may be used, pyrethroid only and pyrethroid-PBO ITNs. The model parameters relating to ITN impacts are given in Table S6 and a description of how ITN impacts are modelled is below.

**Table S6.** Model parameters used in the 'set_bednets' function of the 'malariasimulation' model. The levels of pyrethroid resistance in the vector population are estimated using the annual predictive maps developed by Hancock et al. [13].

| Parameter | Definition | Value | Source |
|---|---|---|---|
| $d_{N0}$ | Probability of dying during a single feeding attempt | Depends on type of bednet and the level of insecticide resistance, $R_a^y$. See Table S1. | [40] |
| $r_{N0}$ | Probability of repeating during a single feeding attempt | Depends on type of bednet and the level of insecticide | [40] |

| | | | |
|---|---|---|---|
| | | resistance, $R_a^y$. See Table S1. | |
| $r_{Nm}$ | Minimum probability of repeating during a single feeding attempt | 0.24 | [40] |
| $\gamma_N$ | Half-life of insecticidal activity in years | Depends on type of bednet and the level of insecticide resistance, $R_a^y$. See Table S1. | [40] |

The model of the implementation of ITNs used in 'malariasimulation' is described fully in Sherrard-Smith et al. [40] and a summary is provided here. When a person is using a net indoors, we consider that any given Anopheles mosquito feeding attempt can result in either a mosquito being deterred before entering the house or those that enter the house either dying, successful blood-feeding or exiting unfed. These outcomes are summarized as the probabilities of repeating (being deterred or exiting unfed; $r_{N0}$), dying ($d_{N0}$) or feeding successfully (1 - $r_{N0} - d_{N0}$). They depend on the level of insecticide resistance in the vector population, the decay in insecticidal activity following net distribution and the type of bednet used. The dependence on the level of pyrethroid resistance, *R*, is incorporated by a non-linear relationship between the probability of surviving exposure to a pyrethroid-only ITN in an experimental hut trial:

$$1 - l_1 = 1/(1 + \frac{R}{\alpha_1})^{-\alpha_2^2}$$

( 23 )

where $l_1$ is the probability of dying following ITN exposure (across all mosquitoes that enter a hut), *R* is the probability of dying in a WHO standard pyrethroid susceptibility bioassay and $\alpha_1$and $\alpha_2$ are constants. The mortality following exposure to a pyrethroid-PBO ITN, $l_2$, is also dependent on *R* via the following non-linear relationship:

$$l_2 = \frac{1}{1 + \exp\,(-(\beta_1 + \beta_2 l_1)}$$

( 24 )

where $\beta_1$and $\beta_2$ are constants. The probabilities $r_{N0}$ and $d_{N0}$ are simple parametric functions of $l_1$ (for pyrethroid-only nets) and $l_2$ (for pyrethroid-PBO nets), and other constant parameters including the probability that mosquitoes are not deterred by the presence of a net, *m*, and the probability that mosquitoes enter a house and feed in the absence of an intervention, $k_0$ [40].

We use the estimates of $d_0$ and $r_0$ for each net type across different levels of pyrethroid resistance developed by Sherrard-Smith et al. [40], where resistance is measured by standard susceptibility tests. For each area *a*, we estimate the average level of pyrethroid resistance in the vector population in each year *k*, $R_a^y$, as:

$$R_a^y = \frac{1}{P}\sum_{p=1}^{P} R_p^y$$

( 25 )

where $R_p^y$ is the resistance to the pyrethroid deltamethrin in pixel *p* within area *a* in year *y* predicted by the annual geospatial maps of phenotypic resistance to deltamethrin in mosquito species from the *An. gambiae* complex in Sub-Saharan Africa developed by Hancock et al. 2020 [13] (Table S1), and *P* is the total number of pixels in area *a*.

Insecticidal activity wanes after net distribution. The decay rate of the killing activity of the active ingredient over time, $\gamma_N$ again depends on the mortalities following exposure to each type of net, $l_1$ and $l_2$:

$$\gamma_N = \mu_N + \rho_N(l_1 - \tau)$$

where $\mu_N$ and $\rho_N$ are constants that take on different values depending on the type of net, *N* (1 = pyrethroid-only or 2=pyrethroid-PBO) and $\tau$ is a constant.

Data to estimate separate values of $R_p^y$, $d_0$ and $r_0$ for different malaria vector species are unavailable, so we use the same values of these parameters for all vector species groups modelled. Different vector species do, however, experience different rates of mortality from ITNs and IRS due to differences between the species in parameters specifying preferences for indoor blood feeding (rather than biting outdoors) and blood feeding on humans (rather than animals) (Tables S4 & S8).

##### 4.4.2 Bednet distributions

The number of new bednets distributed in each area per year is calculated using the 'netz' package in R [44] using the code available on Github [45]. The 'fit_usage' function in 'netz' was used to estimate the annual number of netz distributed to match to the net usage, defined as the proportion of the human population sleeping under a bednet, in each year of the period 2000-2018. Annual net usage for each area was estimated using publish predictive maps [11] (Tables S1 & S2). The model takes into account loss of nets from the human population due to nets being discarded (i.e. thrown away), and assumes a constant rate of net loss per year. We assume that the duration of bednet retention has a half life of 5 years.

### S4.5 Indoor residual spraying

Our model of the impacts of IRS on rates of mortality and human blood feeding in the vector population is implemented using the 'set_spraying' function in 'malariasimulation' and is provided on [cite Github]. The parameters that we use in the 'set_spraying' function are given in Table S7. The 'malariasimulation' model of IRS impacts is based on the models developed by Sherrard-Smith et al. [46], and is summarised briefly below.

**Table S7.** Model parameters used in the 'set_spraying' function of the 'malariasimulation' model. The levels of pyrethroid resistance in the vector population are estimated using the annual predictive maps developed by Hancock et al. [13].

| Parameter | Definition | Value | Source |
|---|---|---|---|
| $l_{s\theta}$ | The proportion of indoor blood-feeding mosquitos that are killed in a house immediately after spraying | Depends on the insecticide product and the levels of resistance; see Table S1. | [46] |
| $l_{s\gamma}$ | Determines the rate of decay mosquito | Depends on the insecticide product | [46] |

| | | | |
|---|---|---|---|
| | mortality due to IRS in days since spraying | and the levels of resistance; see Table S1. | |
| $k_{s\theta}$ | The proportion of indoor blood-feeding mosquitos successfully blood-feed in a house immediately after spraying | Depends on the insecticide product and the levels of resistance; see Table S1. | [46] |
| $k_{s\gamma}$ | Determines the rate of increase in rates of successful blood feeding with days since spraying | Depends on the insecticide product and the levels of resistance; see Table S1. | [46] |
| $m_{s\theta}$ | The proportion of mosquitoes deterred from entering a sprayed house immediately after spraying | Depends on the insecticide product and the levels of resistance; see Table S1. | [46] |
| $m_{s\gamma}$ | Determines the rate of decay in deterrence with days since spraying | Depends on the insecticide product and the levels of resistance; see Table S1. | [46] |

IRS is assumed to impact the proportion of indoor-feeding mosquitoes (Table S8) that are killed ($l_s$), successfully fed ($k_s$), or deterred away ($m_s$) from a sprayed house. These probabilities depend on the insecticide product and the time since spraying. For pyrethroid sprays, they also depend on the level of pyrethroid resistance in the vector population.

Each of these probabilities depends on time since spraying according to the following parametric functions:

$$l_s = \frac{1}{1 + \exp\left(-(l_{s\theta} + l_{s\gamma} t)\right)}$$

$$k_s = \frac{k_0}{1 + \exp\left(-(k_{s\theta} + k_{s\gamma} t)\right)}$$

$$m_s = \frac{1}{1 + \exp\left(-(m_{s\theta} + m_{s\gamma} t)\right)}$$

*( 26 )*

where $l_{s\theta}$ , $k_{s\theta}$ and $m_{s\theta}$ are values at the time of spraying and $l_{s\gamma}$, $k_{s\gamma}$ and $m_{s\gamma}$ are impact durations.

The association between mosquito mortality measured in WHO standard susceptibility tests and mortality induced by IRS in experimental huts is also used to estimate the diminishing impact of pyrethroid-based IRS with resistance. Sherrard-Smith et al. [46] developed parametric functions similar in form to eqn (23) describing how the values at the time of spraying, $l_{s\theta}$ , $k_{s\theta}$ and $m_{s\theta}$, and the rates of decay $l_{s\gamma}$, $k_{s\gamma}$ and $m_{s\gamma}$ are affected by pyrethroid resistance $R_a^k$ (Table S1). We use the values of pyrethroid resistance in each area for each year, $R_a^y$, in our parameterisation of pyrethroid IRS effects, assuming that resistance levels are the same for all types of pyrethroid insecticides. We assume that mosquitoes had no resistance to organophosphate or carbamate insecticides. Experimental hut data to parameterise effects of organochlorine IRS insecticides is not available, so we assume that effects of organochlorines are the same as those of pyrethroids. Data describing IRS efficacy for different vector species are unavailable, so we use the same IRS parameterisation for all vector species modelled.

### S4.6 Vector interactions with insecticidal interventions

Vector biting behaviour, and how this interacts with the presence of ITNs and IRS, is modelled as described in Griffin et al. [23], who follows the approach taken by Le Menach et al. [47]. This model is parameterised in 'malariasimulation' using the 'set_species' function. The parameter values that we use in 'set_species' are given in Table S8.

**Table S8.** Vector behaviour parameters used in models of the impacts of ITNs and IRS

| **Parameter** | **Description** | ***An. gambiae* and *An. coluzzii*** | ***An. arabiensis*** | ***An. funestus*** | **Source** |
|---|---|---|---|---|---|
| $\Phi_I$ | Proportion of bites that are taken indoors | 0.97 | 0.96 | 0.98 | [40] (within the maximum and minimum values reported) |
| $\Phi_B$ | Proportion of bites that are taken indoors and bed | 0.89 | 0.9 | 0.9 | [40] (within the maximum and minimum values reported) |
| $k_0$ | The proportion of mosquitoes that enter a house and successfully feed and exit | 0.69 | 0.69 | 0.69 | [23,40] |

### S4.7 RTS,S vaccines

We model implementation of RTS,S vaccination based on the recommendation from the World Health Organisation that countries should implement a four-dose schedule in children from five months of age [48]. We use the 'set_rtss_epi' function in 'malariasimulation' to specify that children who reach five months of age are given a three-dose primary series and a booster dose at the same time in the following year. We assume a vaccination coverage of 80%, meaning that 80% of children will receive the primary dose series upon reaching five years of age, and a booster coverage of 64% to represent a drop-off in uptake of boosters. We use the default parameterisation given in 'malariasimulation' of vaccine efficacy, and how this decays over time (Table S9), which is based on the vaccine antibody model developed by Hogan et al.[49].

**Table S9.** Parameters of the model of RTS,S vaccine antibody model

| Parameter | Definition | Value | Source |
|---|---|---|---|
| $V_{max}$ | Maximum efficacy against infection | 0.93 | [49,50] |
| $\alpha$ | Dose response shape parameter | 0.74 | [49,50] |
| $\beta$ | Dose response scale parameter | 99.4 EU/mL | [49,50] |
| $CSP_{peak}$ | Peak anti-CSP following the third dose | 621 (geometric mean, EU/mL) | [49,50] |
| $CSP_{fourth}$ | Peak anti-CSP following the fourth dose | 277 (mean on log-scale) | [49,50] |
| $\rho_{peak}$ | Proportion of short-lived response following primary schedule | 0.88 | [49,50] |
| $\rho_{fourth}$ | Proportion of short-lived response following the fourth dose | 0.7 | [49,50] |
| $1/r_S$ | Half-life of short-lived antibodies | 45 (mean in days) | [49,50] |
| $1/r_l$ | Half-life of long-lived antibodies | 591 (days, sampled from log-normal distribution) | [49,50] |

#### S4.8 Relationship between malaria transmission and prevalence

Malaria transmission rates vary across the modelled areas due to variation in the vector population abundance and species composition, and the coverage of insecticidal vector control interventions (ITNs and IRS) in the human population. The relationship between malaria prevalence in the year immediately prior to the first release of gene drives and the ratio of the total vector population size to the total human population size in each area is non-linear, with a higher gradient occurring at lower values of the number of mosquitoes per human (Figure S2). This pattern is what we would expect to see based on the known relationship between malaria transmission and prevalence [51].

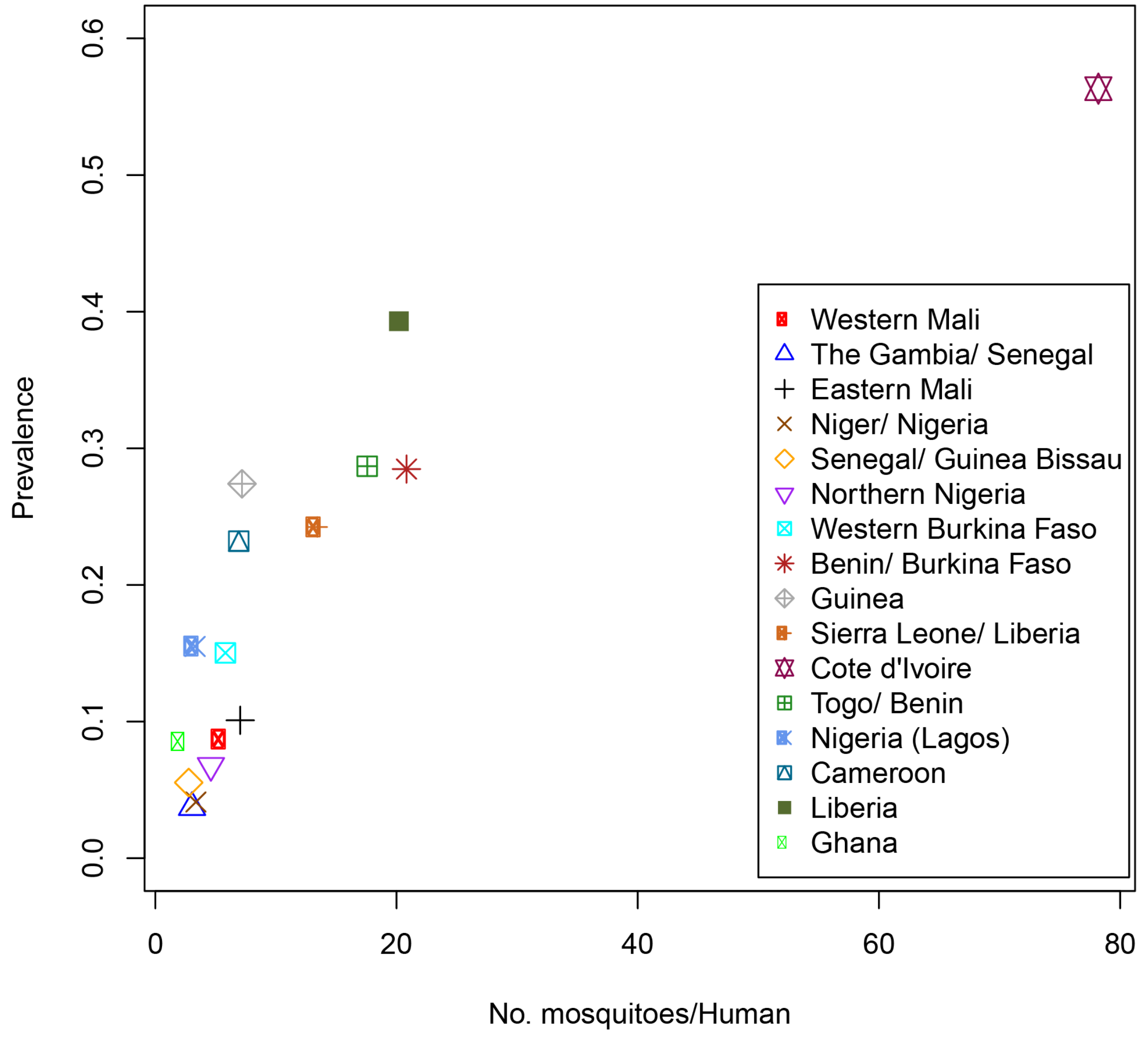

**Figure S2.** The average annual prevalence of *P. falciparum* malaria predicted by the model vs the average annual total number of vectors per human resident, for each modelled area. Data show predictions from the malaria transmission dynamic model for the year immediately prior to the first releases of gene drives.

### S5. Gene drive frequencies in each square 12 years post-release

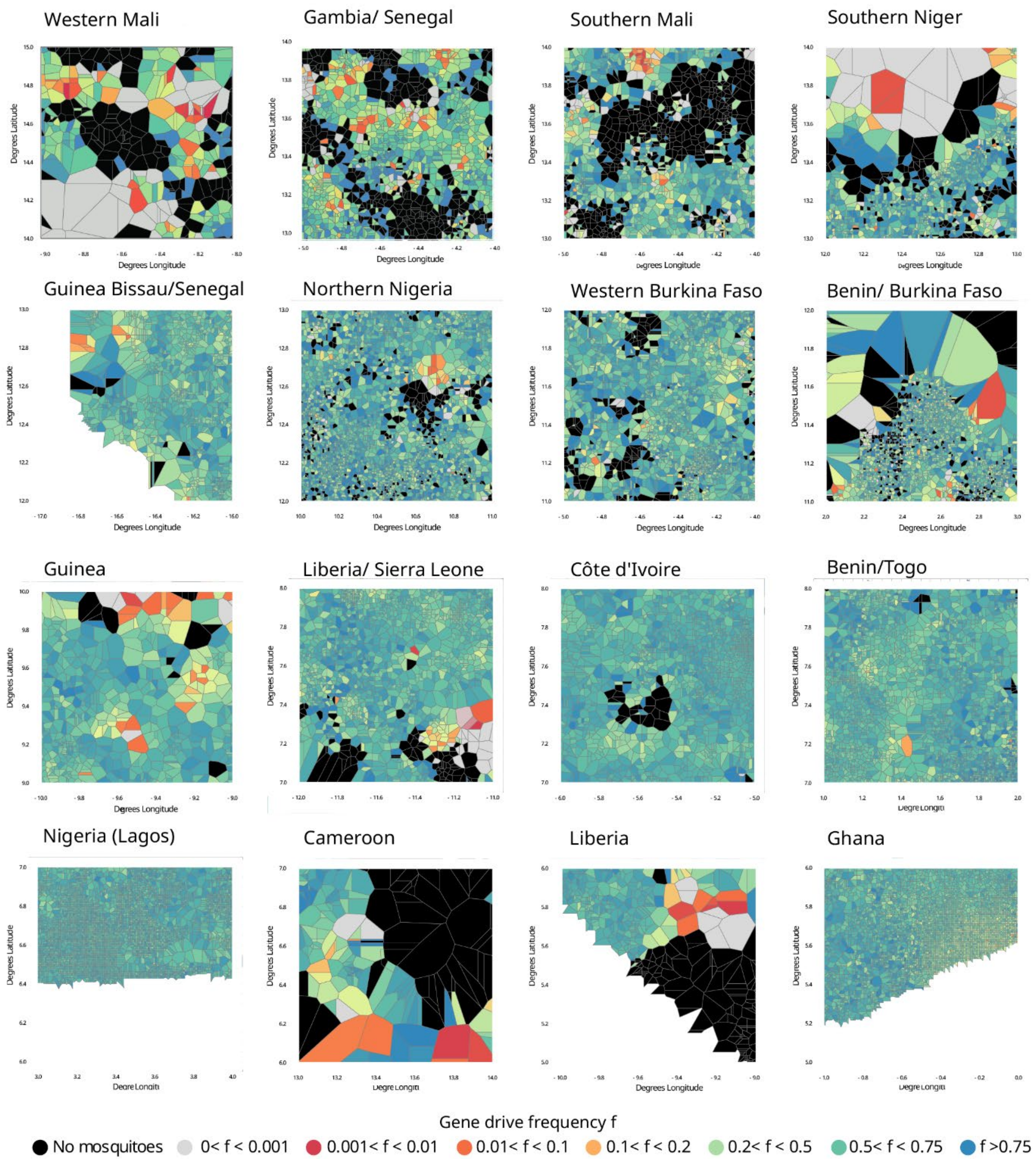

**Figure S3.** Snapshots of typical simulations of gene drive releases in each of the study areas. In each plot, the localities are coloured by gene drive frequency 12 years after releases in 50 random localities.

### S6. Sensitivity to $R_m$

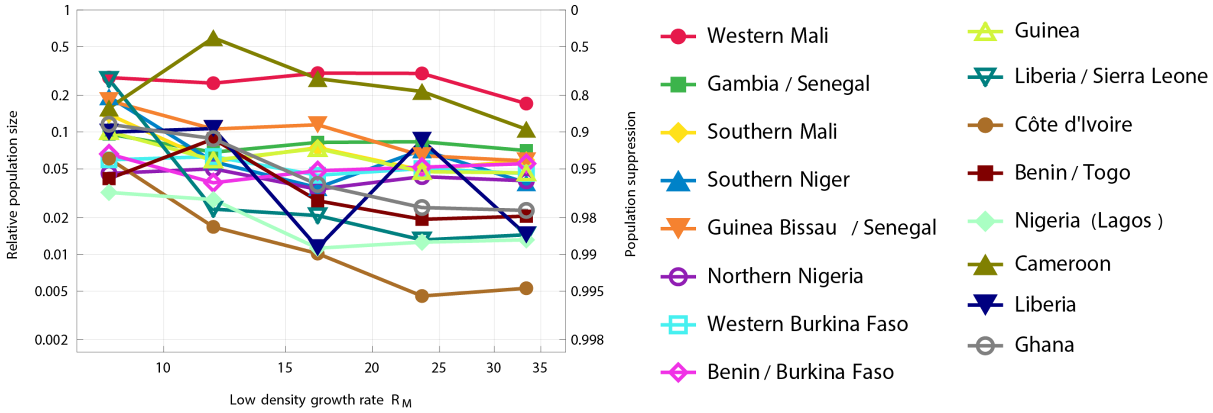

**Figure S4.** Sensitivity of population suppression to the maximum population growth rate $R_m$. Each marker shows the predicted suppression of the entomological model 12 years after gene drive release, from a single simulation run. All parameters are at default settings, except the density independent juvenile mortality, $\mu_{JUV}$, which is altered to manipulate $R_m$, and the area specific carrying capacity parameters $\{K_{1a}, K_{2a}\}$ which are altered such that the area-specific pre-gene drive population sizes were unchanged. $R_m$ is calculated in this figure as the product: (juvenile survival probability at low density) x (average adult longevity) x (adult female egg laying rate) x (½), where the latter factor of a half is because half of eggs are female.
